# Metabolic control of mitochondrial plasticity and extracellular vesicle biology drives Cryptococcus neoformans virulence

**DOI:** 10.64898/2026.03.02.708673

**Authors:** Nan Hong, Xuexin Bai, Hongshan Yuan, Pan Yu, Hannah Edwards, YiNing Ma, Wanqing Liao, Hulin Chen, Qi Zheng, Yang Wang, MengMeng Wang, Jianping Xu, Min Chen, Matthew Fisher

**Affiliations:** Department of Dermatology, Jinling Hospital, Affiliated Hospital of Medical School, Nanjing University, Nanjing, China; Department of Dermatology, Shanghai University of Medicine & Health Sciences Affiliated Zhoupu Hospital, Shanghai, China; Department of Infectious Disease Epidemiology, MRC Centre for Global Infectious Disease Analysis, School of Public Health, Imperial College, London, UK; Department of Burns and Plastic Surgery, Jinling Hospital, Affiliated Hospital of Medical School, Nanjing University, Nanjing, China; Shanghai Key Laboratory of Molecular Medical Mycology, Changzheng Hospital, Shanghai, China; Department of Dermatology, Guangdong Women and Children Hospital, Guangzhou, China; Center of Molecular Metabolism, Nanjing University of Science and Technology, Nanjing, China; Kidney Intensive Care Unit, National Clinical Research Centre of Kidney Diseases, Jinling Hospital, Affiliated Hospital of Medical School, Nanjing University, Nanjing, China; Clinical Laboratory Medicine Center, Yueyang Hospital of Integrated Traditional Chinese and Western Medicine, Shanghai University of Traditional Chinese Medicine, Shanghai, China; Department of Biology, McMaster University, Hamilton, Canada

**Author notes:** ***Corresponding authors:*** Matthew Fisher, Department of Infectious Disease Epidemiology, School of Public Health, Imperial College, 80 Wood Lane, London W12 0BZ, United Kingdom.; Min Chen, Department of Dermatology, Shanghai University of Medicine & Health Sciences Affiliated Zhoupu Hospital, Zhouyuan Road 1500, Shanghai, China.; Nan Hong, Department of Dermatology, Jinling Hospital, 305 Zhongshan Road, Nanjing, China. Contributed equally (Nan Hong, Xuexin Bai, Hongshan Yuan, Pan Yu).

**Keywords:** *Cryptococcus neoformans*, VNIa-5 subclade, Extracellular vesicles, Mitochondrial morphology, Glucose

## Abstract

Metabolic adaptation to nutrient stress is a key but poorly understood driver of fungal virulence. Here, we show that a dominant East Asian lineage of *Cryptococcus neoformans* (VNIa-5), which disproportionately infects immunocompetent hosts, has undergone lineage-specific rewiring of glucose-responsive stress pathways. Integrating population genomics, transcriptomics, and experimental infection models, we demonstrate that VNIa-5’s clinical dominance is not explained by environmental prevalence. Instead, selective activation of Snf1 signalling links glucose limitation to mitochondrial tubularisation, extracellular vesicle remodelling, and enhanced melanization. Under low-glucose conditions, VNIa-5 exhibits marked mitochondrial plasticity and extracellular vesicle compositional shifts resembling hypervirulent outbreak lineages of *Cryptococcus gattii*. Following experimental induction of dormancy, VNIa-5 shows significantly increased virulence *in vivo* compared with the closely related but clinically rare VNIa-31 subclade, with host survival tightly correlated with mitochondrial morphology. These findings identify metabolic stress integration as a central mechanism shaping cryptococcal virulence and disease in immunocompetent human hosts.

## Introduction

Cryptococcosis is a life-threatening fungal disease that affects people worldwide. The basidiomycete yeast *Cryptococcus neoformans* – comprising the VNI/VNII/VNB lineages - accounts for the vast majority of cryptococcal meningoencephalitis (CM) cases globally and is associated with a high mortality^1^, leading to its recent inclusion in the World Health Organization fungal priority pathogen list. Among these lineages of *Cryptococcus neoformans*, the globally distributed VNI and the rarely detected VNII are predominately clonal^2^, whereas VNB exhibits frequent recombination and is geographically restricted to sub-Saharan Africa and South America^3–5^. In Asia, *C. neoformans* VNI is the dominant lineage and is thought to have undergone a relatively recent and rapid population expansion, resulting in low overall genetic diversity structured into three dominant phylogenetic subclades: VNIa-4, VNIa-5 and VNIa-93^6^.

Across temperate, subtropical and tropical regions of Asia, *C. neoformans* VNIa is commonly recovered from avian excreta and decayed wood within tree-trunks^7–9^. The capacity of *C. neoformans* to cause disease is widely considered an incidental consequence of environmental adaptation, shaped by exposure to diverse ecological stressors and microbial predators^10^. However, epidemiological and evolutionary studies of *C. neoformans* in Asia have largely relied on low-resolution genotyping approaches, such as multi-locus sequence typing (MLST) and amplified fragment length polymorphism (AFLP), which lack the resolution needed to link genomic variation to virulent phenotypes. This limitation is exemplified by the discovery that the hypervirulent MLST ST93 strain comprises two genomically distinct clades (ST93A and ST93B) with markedly different virulence profiles^11^. More broadly, conflicting results from phenotypic assays across subclades^12^ highlighting that the biological determinants of virulence and host risk remain poorly resolved in this highly populated region.

One subclade of particular importance is *C. neoformans* VNIa-5 (previously named as VNI-γ or sequence type 5 – ST5) which is widely distributed across China, Japan, Korea, and Vietnam and is strongly associated with non-HIV cryptococcal infections^13–17^. In contrast to sub-Saharan Africa where the dominant risk factor is infection with HIV, non-HIV and immunocompetent patients are increasingly at risk of infection in Asia and account for 20%–50% of the total cryptococcosis cases^18,19^. Further, it has been reported that cryptococcosis in non-HIV patients is associated with worse outcomes than those in AIDS patients^20^. China is among the most heavily impacted countries for non-HIV cryptococcosis, with an estimate of 65,507 novel CM cases annually, corresponding to an incidence of 4.57 per 100,000 persons per year^21^.

Strikingly, in China more than 80% of HIV-uninfected CM patients are infected by VNIa-5^22–29^, a pattern that is mirrored in Vietnam and South Korea^14,30^. Although VNIa-5 has been isolated from clinical specimens outside Asia-including Africa, Europe, and North America^6^, its global clinical prominence contrasts sharply with its low environmental prevalence, which typically ranges from 6 – 20%^7,8^. This discordance raises important questions central to host-pathogen biology: why does VNIa-5 cause a disproportionate burden of disease despite its limited environmental representation, and what cellular, metabolic, and genomic features distinguish it from co-circulating *C. neoformans* lineages? To address these questions, we conducted an integrated population genomic, transcriptomic, phenotypic, and *in vivo* analysis of clinical and environmental isolates from China to define the molecular mechanisms underpinning the enhanced virulence of VNIa-5.

## Results

### Isolate characteristics

A total of 81 Chinese isolates were obtained and sequenced, including 53 environmental isolates and 28 clinical *C. neoformans* isolates. All clinical isolates were collected from CSF specimens of HIV-negative patients. All 53 environmental isolates were collected from pigeon droppings across eastern China and identified as VNI genotype *C. neoformans*. In this study, we used the Qinling-Huaihe line (roughly 33rd parallel North) to distinguish Northern China (temperate zone) and Southern China (tropical and subtropical zones). Twenty-one environmental isolates were from Northern China and 32 environmental isolates from Southern China. For environmental isolates in China, VNIa-5 accounted for a larger fraction in the temperate zone (38.1%, 8/21) than that in the subtropical / tropical zones (21.87%, 7/32). In contrast, VNIa-31 showed a similar distribution between temperate (52.38%, 11/21) and subtropical / tropical zones (46.87%, 15/32). Population genetic analysis showed environmental isolates in subtropical / tropical zones (π = 0.0014) had slightly higher genetic diversity than environmental isolates from temperate zone (π = 0.0012) (Figure 1A & Table S1).

**Figure 1.**
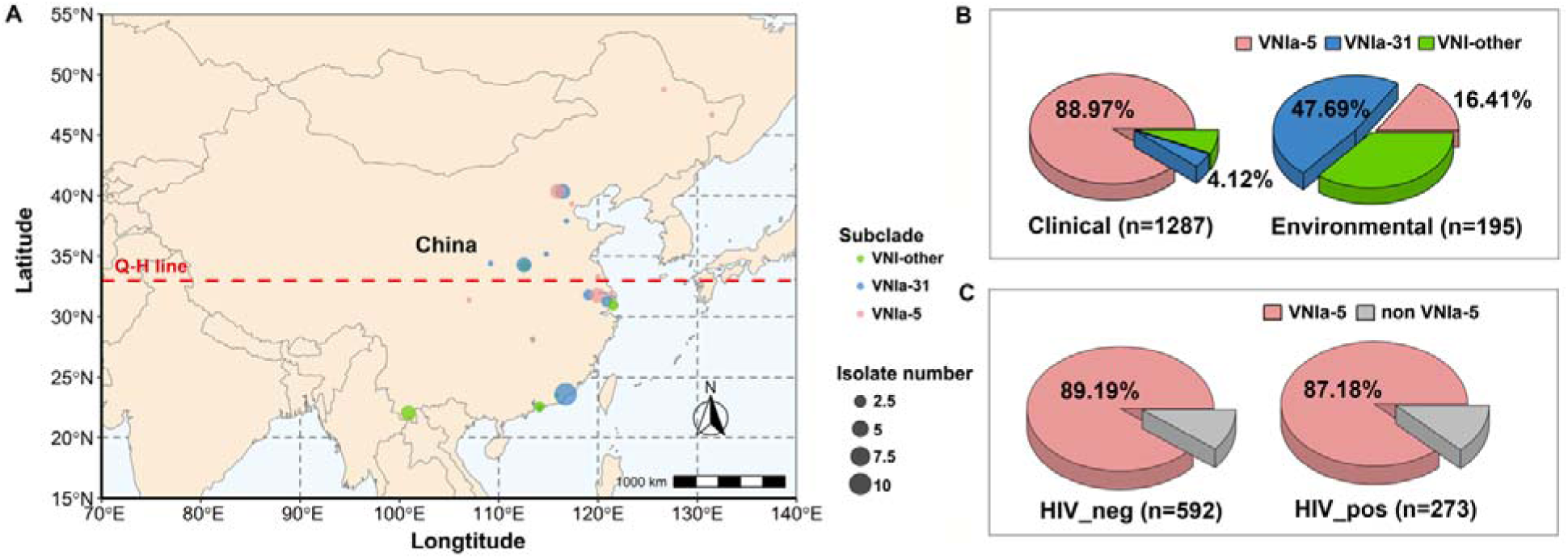
Environmental sampling and pooled MLST analysis of *C. neoformans* isolates in China. (A) All 53 environmental isolates in this study were sampled from pigeon droppings mainly from East China. Q-H line: Qinling-Huaihe line to distinguish Northern and Southern China. (B) MLST analysis based on 1287 clinical isolates and 195 environmental isolates from China showed VNIa-5 to be clinically prevalent but infrequently sampled from environmental specimens. (C) MLST comparison based on 592 non-HIV patients and 273 HIV-infected patients revealed VNIa-5 to dominate in both groups.

### Environmental isolates from China show low-diversity with VNIa-31 dominant

Here we retain the naming scheme of Desjardins et al.^4^ and Ashton et al.^6^ and refer to the subclades within VNI after the predominant MLST sequence type in each subclade. A phylogeny (Figure 2A) based on 81 Chinese isolates and 8 reference strain was constructed based on 155,594 variable sites across cryptococcal genome. Of the 81 Chinese *C. neoformans* isolates presented here, 98.76% were identified as VNIa while only one isolate was VNIb. With VNIb as the outgroup, the phylogeny shows that the population structure of VNIa isolates in China is dominated by two clonal subclades (VNIa-31 and VNIa-5). In addition to the three predominant VNI subclades discovered in Asian clinical isolates (VNIa-5, VNIa-4 and VNIa-93)^6^, VNIa-31 is newly shown in this study as a 4^th^ common subclade, especially amongst environmental samples where it comprised half of the recovered isolates (49.06%, 26/53). Here VNIa-31 clustered with the VNIa-outlier reference strain, which accounted for a fractional proportion of Ashton’s data (1.27%, 11/863)^6^. This likely owes to the small number of environmental isolates (2.89%, 25/863) enrolled in Ashton’s study^6^ alongside its geographical focus regions of southeast Asia outside of China.

**Figure 2.**
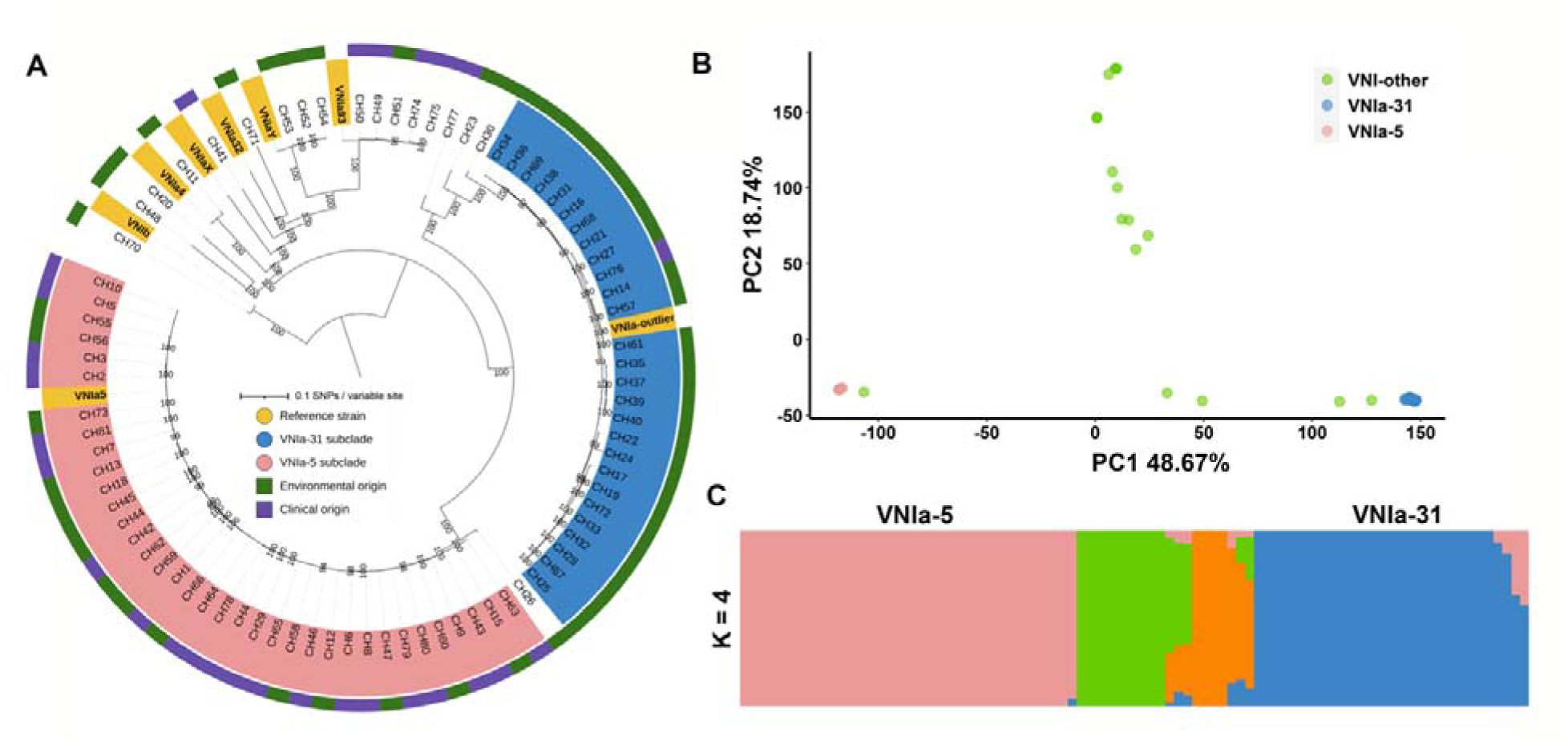
Whole-genome sequencing supports classification of VNIa-5 and VNIa-31 as two distinct and highly clonal sub-clades. (A) Maximum likelihood phylogeny of 89 *C. neoformans* genomes, including 81 Chinese isolates and 8 reference strains from Ashton’s study^6^ representing different VNI subclades. (B) PCA showing VNIa-5 and VNIa-31 to be two clonal subclades with high divergence. (C) Unsupervised ADMIXTURE clustering analysis of all isolates at *K =* 4.

Chinese environmental *C. neoformans* isolates possessed comparatively low genetic diversity (π = 0.0013) when compared to the VNI population mainly from Africa and Europe (π = 0.0021)^4^ as well as VNI population mainly from India and Thailand (π = 0.0020)^3^. VNIa-31 showed high clonality with extremely low genetic diversity (π = 0.00014) and accounted for half of Chinese environmental isolates. VNIa-31 isolates were clustered in a highly clonal subclade with robust phylogenetic support (100% bootstrap), showing little to no intermixing with other phylogenetic groups (Figure 2A). Unsupervised model-based clustering also identified highly structured ancestry components enriched in VNIa-31. The clustering solution with the lowest cross validation error value (*K* = 4) grouped the VNIa-31 isolates into a single genetically homogenous group (Figure 2C) with F_ST_ values of VNIa-31 compared against the remaining three groups all larger than 0.25. Together, these metrics demonstrate that VNIa-31 is a distinct but genetically homogenous group that has diverged from other subclades.

### VNIa-5 predominant in clinical but infrequent in environmental isolates

The VNIa-5 subclade, a major cause of cryptococcal infections in HIV-uninfected patients, has been previously highlighted as the predominant cryptococcal pathogen in east Asia. Here we combined the data from this study and all 12 previously published studies^7,8,22–29,31,32^ from China with available MLST data, resulting in a pooled dataset of 1,287 clinical and 195 environmental *C. neoformans* isolates from China (Figure 1B & Table S1). VNIa-5 predominates in clinical isolates (88.97%,) but is rare in the environment (16.41%). This striking pattern indicates that VNIa-5’s clinical predominance is not caused by frequent environmental exposures but rather owes to its unique virulence characteristics. In stark contrast, VNIa-31 accounts for only 4.12 % of clinical isolates but 47.69% of environmental isolates (Figure 1B). Principle component analysis (PCA) shows both VNIa-5 and VNIa-31 to be highly clonal and diverged from each other with a mean F_ST_ value of 0.63 (Figure 2B).

### Genetic and transcriptomic features of VNIa-5 associated with virulence phenotypes

We next identified regions of the VNIa-5 genome that are associated with selective sweeps, based on metrics of SNP diversity. Regions with high π ratio (comparing between VNIa-5/VNIa-31) identified 11 candidate regions that possess larger number of variants, indicating these regions may be under diversifying selection. Here, candidate regions under selection were defined as outliers falling concurrently within the upper quantile of the distribution of F_ST_ (> 0.75) and log π ratio (>0.5). Figure 3A identifies the upper intersection of F_ST_ and log π ratio where the 11 candidate regions lie. Here, a total of 49 genes were identified in these 11 regions, of which many were involved in stress response, vesicle transport and mitochondrial function (Table S2). Amongst these, the genes *VEP5* and *SUR7* encode proteins that were recently identified in the proteome of cryptococcal extracellular vesicle (EV)^33^. The Sur7 protein has also been identified as EV-protein markers in *Candida albicans* and in the wheat pathogen *Zymoseptoria tritici*^34,35^. Notably, genes *SNF1* and *SIP1* encode essential components of the Snf1 pathway that controls low glucose response and the utilization of alternate carbon sources^36^, and were also identified in these 11 diverged regions (Table S2).

**Figure 3.**
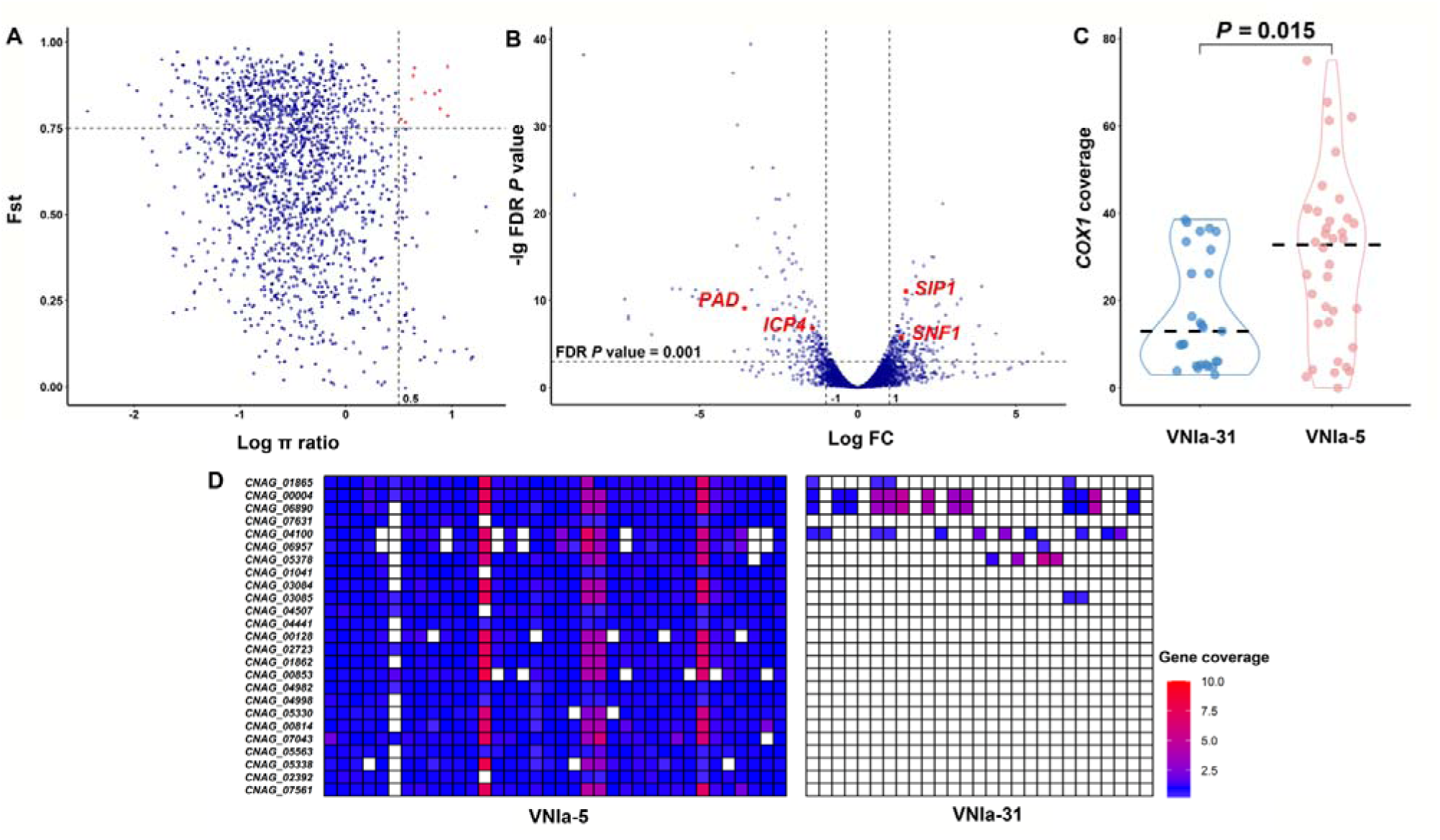
Multi-omics analyses revealed VNIa-5 genomic features concerning EV production, mitochondrial morphology, and low-glucose response. (A) Genome scanning of pairwise F_ST_ combined with π ratio (π VNIa-5/π VNIa-31) revealed 11 regions under selective sweep in VNIa-5. (B) Transcriptional profiling revealed 4 genes (*PAD*, *ICP4*, *SIP1* and *SNF1*) with differential expressions and under selection in VNIa-5. (C) VNIa-5 possess twice *COX1* gene copy number of VNIa-31. (D) CNV analysis revealed 25 genes unique to VNIa-5. Fst: Pairwise F_ST_ value between VNIa-5 & VNIa-31; Log FC: Log_2_(Fold change).

More widely across the genome, when analyzing gene copy number variation, we detected a total of 25 genes that are unique to VNIa-5 when compared against VNIa-31-this included 12 genes with known and 13 genes with unknown functions (Figure 3D & Table S3). Those genes with known function are mainly associated with sugar transportation and metabolisms. Among those genes unique to VNIa-5, the hexose transporter (*CNAG_01862*) is reported to be regulated by Snf1 pathway in exponential growth and mitochondrial respiration of *Saccharomyces cerevisiae*^37^. Moreover, the gene *COX1* is closely related to mitochondrial energy production^38^, and was found to have a median coverage of 32.75 in VNIa-5, twice of that in VNIa-31 with a median *COX1* coverage of 12.98 (*P* = 0.015, Mann-Whitney *U*-test) (Figure 3C). Furthermore, a positive relationship was reported between expression of *COX1* and intra-macrophage proliferation of *C. neoformans*^39^. Thus, the high *COX1* coverage found in VNIa-5 might indicate enhanced intercellular proliferation ability of VNIa-5.

We compared the transcript profiles of VNIa-5 and VNIa-31 growing in YPD broth at 37°C for 24h. A total of 381 protein-coding genes were found to be differentially-expressed genes (DEGs) between VNIa-5 and VNIa-31 with at least 2-fold change and a false discovery rate (FDR) less than 0.001 (Figure 3B & Table S4). Gene Ontology analysis enriched these DEGs into three main aspects: transmembrane transport, membrane component and oxidoreductase activity (Table S4). Among these DEGs, five genes encoding components in the proteome of cryptococcal EVs^33^ included *VEP14*, *RIL1* and *CFO2* were upregulated in VNIa-5, while *EXG104* and *SUC2* were downregulated. Eleven DEGs were found to be functionally related to iron uptake, DNA repair, and RNA processing in mitochondria. Noteworthily, four DEGs (*PAD*, *ICP4*, *SNF1* and *SIP1*) were found to be located in the regions under selection that we identified (Figure 3B & Table 1) -*PAD*, encoding phenolic acid decarboxylase related to yeast utilization of decayed wood, was previously reported to be associated exclusively with the VNIa-5 strains^30^. Of key interest, *SNF1* and *SIP1* are essential components of the Snf1 pathway that mediates the glucose deficiency response, and were both upregulated and under selection in VNIa-5. Together, this implicates low glucose adaptation as a potential virulence characteristic of VNIa-5.

**Table 1.**
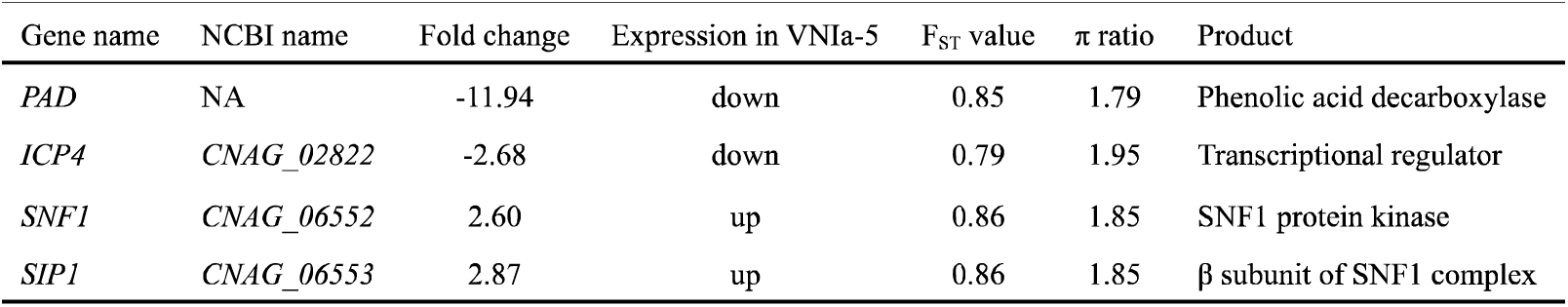
Four genes with differential expression and under selection in VNIa-5. (“NA” Not available; “π ratio” πVNIa-5/ πVNIa-31).

### Active small EV production and enhanced melanin production observed in VNIa-5

The Chinese VNI strains in this study had a broad range of EV sizes, with diameters varying from 15 to 505 nm and a major peak in the 105 nm diameter range (Figure 4 & Table S5). Fungal EVs are broadly classified into two subpopulations based on subcellular origin: small EVs (sEV, exosomes, 30 to 200 nm; non-conventional secretion from endosomes) and large EVs (LEV, microvesicles,100 nm to 1 µm; conventional secretion from plasma membrane)^40–43^. Shown in Figure 4C&D, the sEV fraction (Number of EV_35-195 nm_ / Total number of EVs) was measured across the 81 isolates in our study, showing that VNIa-5 possesses distinctly larger fraction of sEV than VNIa-31, with robust significance (*P* = 1.75E-05, Mann-Whitney *U*-test). In this study, the 28 clinical isolates also showed a higher small EV fraction (*P* = 0.00031, Mann-Whitney *U*-test) than that of the 53 environmental isolates (Table S5), likely due to the predominance of VNIa-5 in clinical isolates.

**Figure 4.**
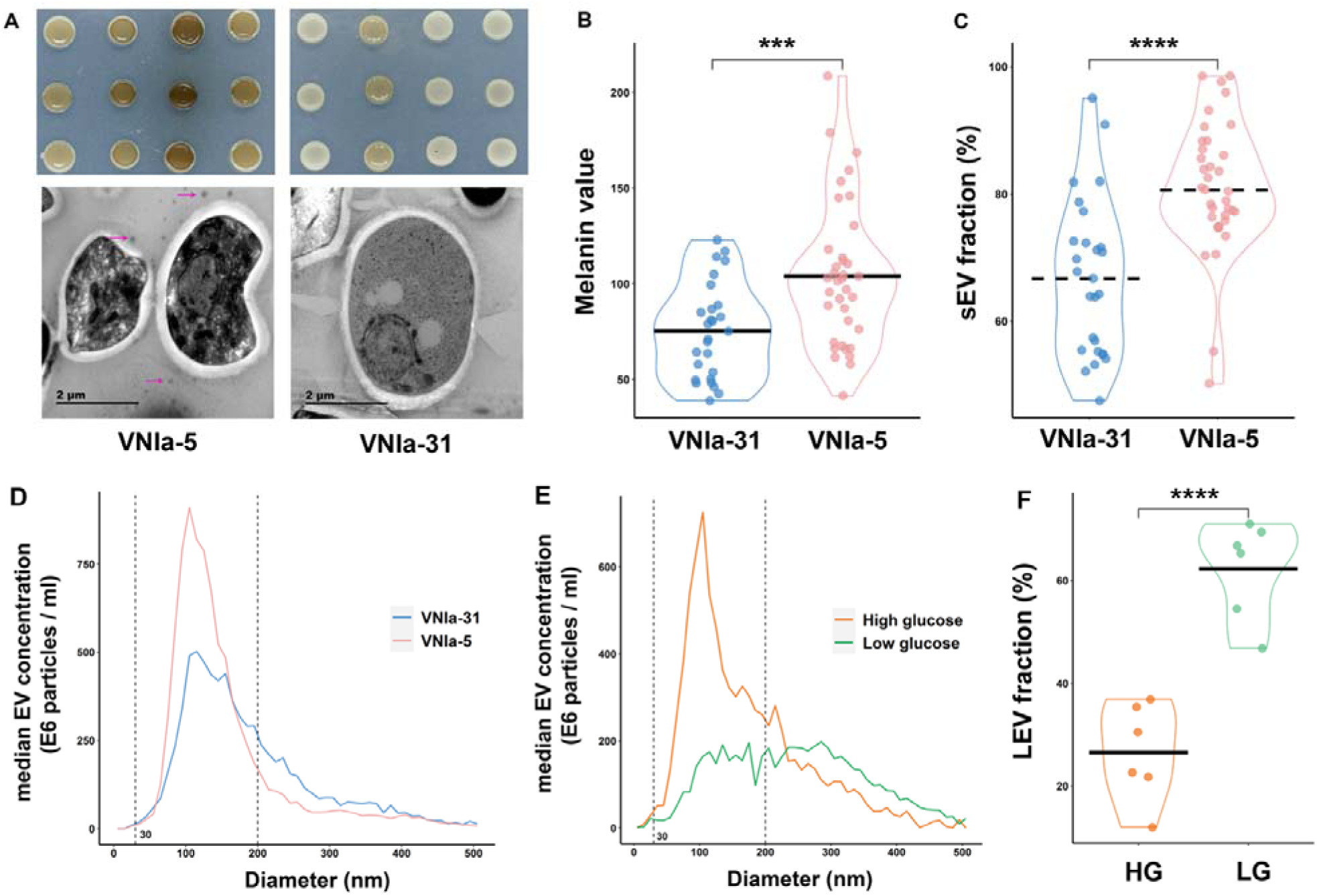
Active sEV production, glucose-responsive EV composition alteration and enhanced melanization observed in VNIa-5. (A) Cryo-electron transmission microscopy showing VNIa-5 produced large number of EVs. Purple arrows point to EVs. VNIa-5 subclade showed more melanin production than VNIa-31. (B) Melanization measurement showed VNIa-5 possess enhanced melanization capacity than VNIa-31. Black line showed average value. (C) VNIa-5 strains showed significantly larger sEV fraction than VNIa-31 on YPD medium. Dashed line showed median value. (D) NTA curves generated using median value of EV concentration of VNIa-5 and VNIa-31 respectively. (E) NTA curves generated using median value of EV concentration of VNIa-5 under high-glucose or low-glucose. (F) In low-glucose condition, VNIa-5 mainly produced LEVs. sEV: small EV; LEV: Large EV; (* *P* < 0.05; ** *P* < 0.01; *** *P* < 0.001; **** *P* < 0.0001).

The unconventional secretary pathway, regulated by endosomal sorting complexes required for transport (ESCRT), controls sEV production^43^, as well as participating in the trafficking of laccase to the cryptococcal cell wall^44^. The active sEV production of VNIa-5 indicated the possibility of more laccases being transported to the cell wall, resulting in enhanced melanin production ability. Accordingly, we measured the melanin production of VNIa-31 and VNIa-5 strains on L-DOPA agar. As shown in Figure 4A&B, VNIa-5 isolates were found to have enhanced melanization when compared against VNIa-31 isolates (average melanin values of 113.26 compared to 75.52; *P* = 0.00056, *t*-test).

### VNIa-5 EV composition alteration is glucose-responsive

According to the population genomic results in our study, essential genes in the Snf1 pathway are under selection in VNIa-5, suggesting adaptation to low glucose environments. To investigate EV response to glucose, we randomly selected six VNIa-5 isolates and grew them on high-glucose (4g/L glucose) medium, and low-glucose (0.5g/L glucose) medium (to mimic the range of glucose concentrations *in vivo*^45^) at 30°C for 24 hours. Shown in Figure 4E & Table S7, we could observe a sharp decrease of small EV fraction under low glucose condition (average sEV fraction: 37.26%,) compared to that on high glucose medium (average sEV fraction: 73.01%). In stark contrast, an increase of large EVs was observed under low glucose condition (average fraction: 62.28%), twice of that grown on high glucose medium (average fraction: 26.51%) (*P* = 6.9E-05, *t*-test) (Figure 4F.)

### VNIa-5 exhibited enhanced tubular mitochondria morphology but failed to show rapid increase in intracellular proliferation

We incubated *C. neoformans* strains under four conditions for observation of mitochondrial morphology: low-glucose (0.5g/L) DMEM, high-glucose (4g/L) DMEM, low-glucose DMEM with 0.7mM H2O2, high-glucose DMEM with 0.7mM H2O2. Cryptococcal mitochondria in our study showed diffuse, tubular, and fragmented morphologies, similar to observations in previous report^46^. Of note, VNIa-5 strain in low-glucose (0.05%) DMEM showed the most evident tubular mitochondria morphology compared to other incubation conditions (Figure 5). Under low glucose condition, VNIa-5 (average fraction: 37.57%) was found to possess a significantly higher fraction of mitochondrial tubularisation than that of VNIa-31 (average fraction: 9.82%) *(P* = 0.00031, Mann-Whitney *U*-test) (Figure 6A).

**Figure 5.**
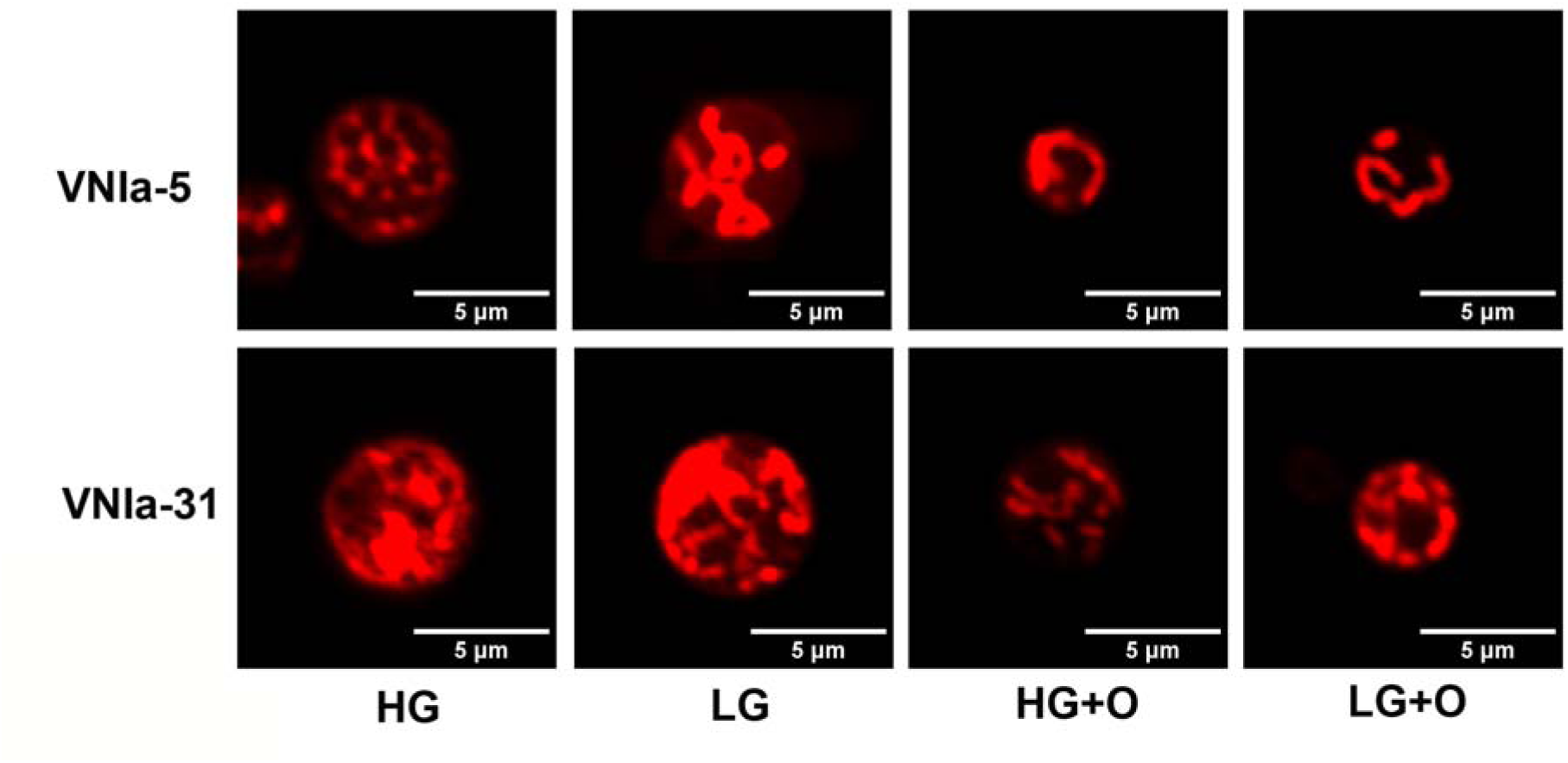
Sample images showing mitochondrial morphology difference of VNIa-5 and VNIa-31. *C. neoformans* mitochondria was stained with MitoTracker CMXRos. Mild oxidative stress was created with 0.7 mM H_2_O_2_ in DMEM. Compared to VNIa-31, VNIa-5 strains showed distinct tubular mitochondria morphology under low-glucose condition. HG: high glucose; LG: low glucose; HG+O: High glucose with mild oxidative stress; LG+O: low glucose with mild oxidative stress.

**Figure 6.**
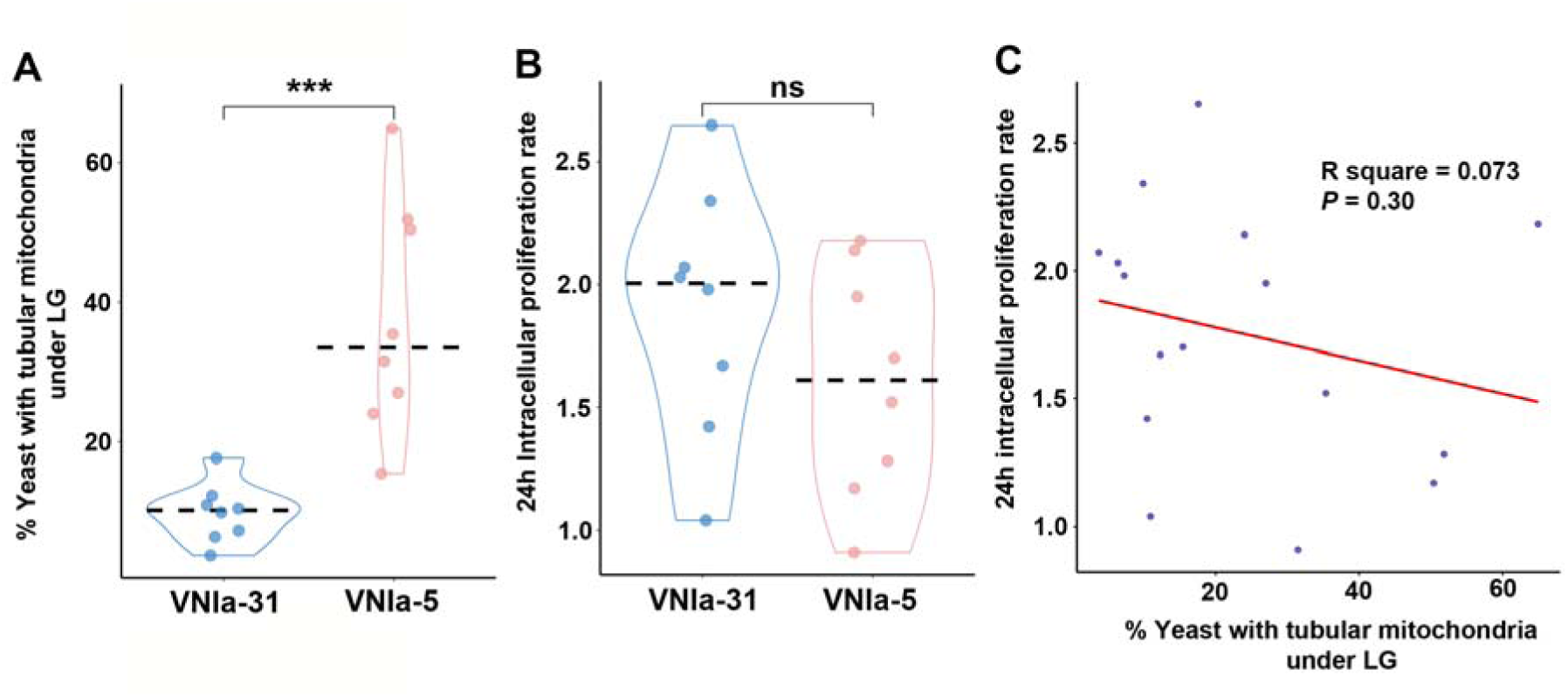
VNIa-5 exhibited enhanced tubular mitochondria morphology but failed to show rapid increase in intracellular proliferation. (A) Under low-glucose condition, VNIa-5 showed distinctly larger fraction of yeasts with mitochondrial tubularisation compared to VNIa-31. (B) VNIa-5 *C. neoformans* showed low IPR 24 h post phagocytosis by J774 cell with an average value of 1.61. (C) In VNI *C. neoformans*, no significant correlation has been observed between mitochondria tubularisation rate and 24 h IPR.

Tubular mitochondria morphology was reported to contribute to intra-macrophage proliferation in Vancouver outbreak *C. gattii* strain^47^. Therefore, we measured the intracellular proliferation rate (IPR) of VNIa-5 strains using J774 cell. However, we observed comparatively low IPR values (average value: 1.60) for VNIa-5 and detected no correlation between IPR and mitochondrial tubularisation in VNIa-5 (R^2^ = 0.0090, *P* = 0.84) within the first 24h post-phagocytosis (Figure 6B&C).

### Dormant VNIa-5 exhibited *in vivo* hypervirulence associated with mitochondrial tubularisation

Consistent with mammals, avian intestines constitute a hypoxic environment^48,49^. This observation may help explain why *C. neoformans* isolates sampled from pigeon droppings and tree trunks experience hypoxic and nutrient-limited conditions, under which *C. neoformans* isolates may switch to a dormant status in order to prolong survival under harsh environment stress^50^. To test this hypothesis, we induced dormancy in VNIa-5 and VNIa-31 cells *in vitro* and then intravenously infected mice (Figure 7A). Strikingly, dormant VNIa-5 (median survival time: 14 days) exhibited hypervirulence, with mice surviving less than half as long as those infected with VNIa-31 (median survival time: 33 days) (Figure 7B). Mouse survival was correlated with the rate of mitochondrial tubularisation under low glucose (Spearman correlation: R^2^ = 0.88, *P* = 0.017) (Figure 7C).

**Figure 7.**
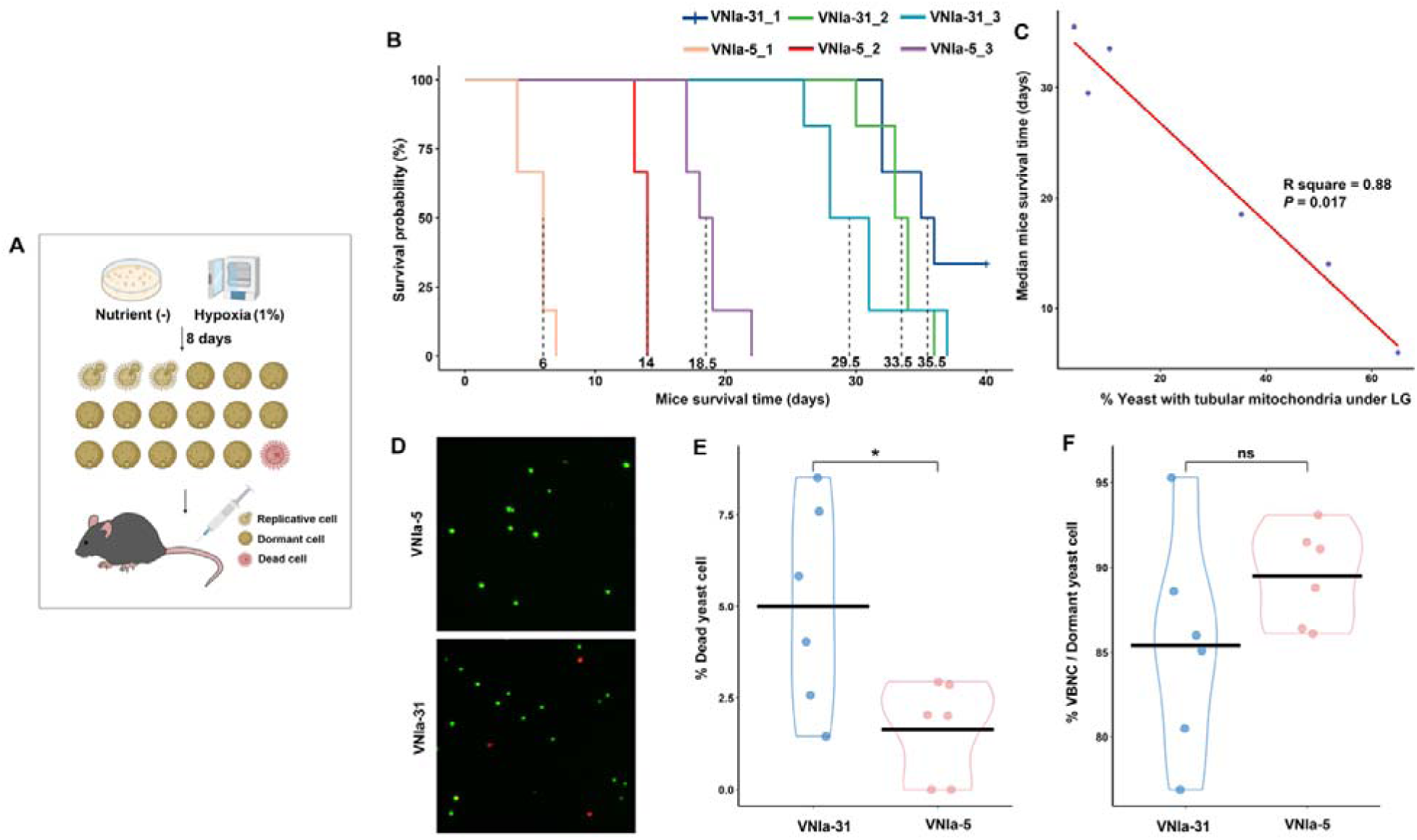
Dormant VNIa-5 exhibited *in vivo* hypervirulence associated with mitochondrial tubularisation. (A) Illustration of dormant VNIa-5 induction process. (B) Dormant VNIa-5 strains showed enhanced virulence compared to dormant VNIa-31 in mice intravenous infection model. (C) Virulence of dormant *C. neoformans* strains correlates with mitochondrial tubularisation. (D & E) Few dead isolates observed by AO/PI staining in VNIa-5 (average: 1.64%) and VNIa-31 (average: 4.99%). (F) After dormancy induction, VNIa-5 and VNIa-31 generated comparable large fraction of VBNCs.

Following dormancy induction, viable-but-non-culturable cells dominated both the VNIa-5 (mean fraction: 91.13%) and VNIa-31 strains (mean fraction: 90.40%) (Figure 7F & Table S7). After *in vitro* dormancy induction, the majority of cells remained viable in both VNIa-5 (average dead cell fraction: 1.64%) and VNIa-31 (average dead cell fraction: 4.99%) (Table S7). Notably, VNIa-31 exhibited a slightly higher dead-cell fraction than VNIa-5 (*P* = 0.032, *t*-test) (Figure 6D&E), indicating that dormant VNIa-5 cells may be more persistent than those of VNIa-31.

## Discussion

Based on the MLST data of 1,287 clinical isolates and 195 environmental isolates in China, our study revealed that VNIa-5 dominates clinical samples (88.97%) but is infrequently detected in the environment (16.41%; Figure 1B). This marked disparity strongly suggests that VNIa-5’s clinical prevalence is driven by intrinsic pathogenic potential rather than environmental abundance. Furthermore, in addition to its dominance among non-HIV cryptococcosis (89.19%, n=592), pooled analyses showed that VNIa-5 also predominates in HIV-infected patients in China (87.18%, n=273) (Figure 1C & Table S1). Given that adaptative immunity is profoundly compromised in HIV-infected patients, the pathogenic advantage of VNIa-5 is likely to arise from differential interactions with host innate immunity.

Environmental *C. neoformans* isolates from China exhibited lower genetic diversity (π = 0.00132) compared with VNI populations sampled predominately from Africa and Europe (π = 0.00209)^4^ and tropical Asia (π = 0.00200)^3^. Notably, our environmental survey identified VNIa-31 as a highly clonal subclade that dominates environmental isolates (47.69%) but is rare in clinical samples (4.12%), and is genomically distinct from VNIa-5 (Figure 2). We further performed multi-omics comparison between VNIa-5 and VNIa-31, and found that VNIa-5 exhibits distinct genetic and phenotypic features, particularly in extracellular vesicle production and mitochondrial morphology. These traits were shown to be glucose-responsive and may plausibly influence cryptococcus interactions with host innate immunity.

Two previous studies have provided insights into the genomic features of VNIa-5, identifying a 165k bp genomic region unique to VNIa-5^30^, and identifying a 21 bp mitochondrial recombinant fragment (with 8 SNPs) unique to this subclade^6^. Notably, in our study the *PAD* gene - located within the previously reported 165k bp genomic region- was found to be both differentially-expressed and under selective sweep (Figure 3). A recent large-scale GWAS research identified nine genes associated with the hypervirulence of VNIa-93 strain^11^, three of which (*CNAG_05664*, *ITR4* and *CNAG_05329*) are involved in amino acid and inositol metabolism: we found these genes were also differentially-expressed between VNIa-5 and VNIa-31 in our study (Table S4). However, no prior study has directly linked the genomic features of VNIa-5 to specific virulence phenotypes. Thanh *et al.* performed phenotype tests but reported that VNIa-5 exhibited similar *in vitro* virulence markers and even hypovirulence in a mouse model^12^, possibly due to their choice of VNIa-4 as a comparator - another prevalent clinical subclade that may share similar pathogenic traits. In contrast, the largely ‘environmental’ VNIa-31, may represent a more suitable comparator.

There has been growing awareness of the importance of fungal EVs in physiology, host-pathogen interactions, and virulence of fungal pathogens. For cryptococcus species, EVs have been shown to modulate host phagocyte activity, promote blood–brain barrier (BBB) invasion in a dose-dependent manner, and transfer virulence traits between strains during infection^41,42^. Our data show that EV composition in VNIa-5 strain is glucose-responsive: under high-glucose conditions, small EVs predominate (average: 73.01%), whereas under low-glucose conditions, large EVs dominate (average: 62.28%) — consistent with recent findings in strain B3501 cultured in rich nutrition versus poor nutrition media^51^. A recent study found *C. neoformans* responds to carbon starvation by increasing capsule porosity and cell wall permeability^52^, which might facilitate LEV secretion. VNIa-5 strains mainly produced large EVs in low-glucose condition (Figure 4E&F), which exhibited enhanced inflammation stimulation ability favoring phagocytosis of cryptococcus^51^. Intravenous treatment of EVs generated under poor-nutrient condition increased the crossing of the BBB by *C. neoformans* and the cerebral fungal load, which reinforces the importance of LEVs in the process of migration and colonization of the fungus in brain tissue^53^. In *S. cerevisiae*, small EVs regulate the release of extracellular gluconeogenic enzymes in response to glucose, facilitating rapid environmental adaptation^54^. Cryptococcal EVs produced in rich-nutrient condition were demonstrated to have inhibitory effects on host innate immune system in both *in vitro* and *in vivo* models^51^. In our study, under high-glucose condition (YPD medium), VNIa-5 produced a significantly greater fraction of sEVs than VNIa-31 (Figure 4C), suggesting enhanced immune evasion via small EV mediated suppression, potentially contributing to its clinical success in immunocompetent hosts.

Fungal melanins mask the fungal cell wall to protect against oxidative stress and laccase is a key virulence factor involved in melanin synthesis. Recently, melanin was found to play an impressive role in dysregulating essential functions of the innate immune system to circumvent host antifungal immunity and promote fungal virulence^55^. Small EVs arise via fusion of multivesicular bodies (MVBs) with the plasma membrane, mediated by the non-conventional pathway that regulated by ESCRT complex^40^. The ESCRT complex was reported to be vital in transporting laccase to the *C. neoformans* cell wall^44^. Therefore, it’s reasonable to hypothesize that VNIa-5 might possess enhanced melanization capacity due to its more active non-conventional secretion. Phenotypic testing supported this, revealing that VNIa-5 produced significantly more melanin than VNIa-31 (Figure 4 A&B), indicating VNIa-5’s advantage in acquiring prolonged intra-macrophage residency.

Carbon source is a key determinant of yeast mitochondrial morphology, which in turn underpins fundamental fungal biological processes^56^. In *Saccharomyces cerevisiae*, carbon availability dictates metabolic strategy (respiration or fermentation) and is associated with distinct mitochondrial morphologies, commonly described as “tubular” or “ring-like”^56^. In *C. neoformans*, prolonged glucose deprivation and hypoxia (8 days) induce dormant subpopulations with viable-but-non-culturable phenotype, accompanied by pronounced mitochondrial remodeling, including increased mitochondrial mass and depolarization^50^. Similarly, in hypervirulent Vancouver Island outbreak (VIO) strains, oxidative stress triggers mitochondrial fusion into a tubular morphology that enhances oxidative stress resistance, while concurrently promoting rapid proliferation of neighbouring, non-tubularized cells within macrophages^47^.

Notably, VNIa-5 isolates exhibited a comparable response, with a high proportion of cells displaying tubular mitochondria under low glucose condition (Figure 5 & Table S6). Quantitatively, VNIa-5 showed a substantially greater fraction of mitochondrial tubularisation (mean fraction: 37.57%) than VNIa-31 (mean fraction: 9.82%) *(P* < 0.001) (Figure 6A). However, despite this mitochondrial phenotype, VNIa-5 strains did not exhibit enhanced early intracellular proliferation within the first 24 hours post-phagocytosis (Figure 6B&C). This result is consistent with prior work, in which intracellular proliferation rate (IPR) assays performed across 14 *C. neoformans* isolates revealed no association between mitochondrial morphology and early intra-macrophage proliferation^47^.

*Cryptococcus*-preferred niches, such as avian intestines and decaying wood within tree trunks, are generally hypoxic and nutrient-limited environment^48^. Under prolonged hypoxia and nutrient-limitation (8 days) *C. neoformans* has been shown to switch to a viable-but-non-culturable or dormant state^50^, suggesting that dormancy may represent a frequent physiological state upon exposure to host-like conditions. To test the impact of dormancy on strain virulence, we induced dormant *C. neoformans* cells *in vitro* and intravenously infected C57 mice. Strikingly, dormant VNIa-5 strains exhibited markedly greater *in vivo* virulence, with a median mouse survival time of 14 days, compared with 33 days for VNIa-31 strains (Figure 7B). Moreover, mouse survival was strongly correlated with the rate of mitochondrial tubularisation rate under low glucose conditions (Spearman correlation: R^2^ = 0.88, *P* = 0.017) (Figure 7C).

Despite a large fraction of VNIa-5 cells exhibiting tubular mitochondria, these strains did not display an immediate increase in IPR, yet nevertheless showed superior *in vivo* virulence that closely correlated with mitochondrial tubularisation (Figure 6&7). One plausible explanation is that *C. neoformans* remain in a dormant state during the early stages of infection to withstand the hostile host environment, followed by reactivation in response to specific *in vivo* stimuli that enable intracellular proliferation. This interpretation is consistent with the chronic disease course observed in most cases of cryptococcal meningoencephalitis ^57^. In support of this model, it has been reported that as early as 24 hours after systemic infection, the majority of *C. neoformans* cells traverse the blood-brain barrier and are engulfed by, or located adjacent to, microglia^58^ which provide an early intracellular reservoir during cryptococcal meningitis^59^. Furthermore, recent work has shown that host innate immune cells can reactivate dormant *C. neoformans* through the secretion of extracellular vesicles and via non-lytic exocytosis^60^. Collectively, these findings support a model in which microglia act as a reservoir that facilitates the persistence and reactivation of dormant *C. neoformans* during infection.

Multi-omics analyses revealed that the Snf1 pathway—a central regulator of yeast adaptation to low-glucose environments—is under selective sweep and transcriptionally upregulated in the VNIa-5 subclade (Figure 3), consistent with an adaptive advantage under glucose-limited conditions. This pathway has also been implicated in mitochondria and extracellular vesicle regulation, including mitigation of Ras-associated mitochondrial dysfunction^61^ and glucose-dependent modulation of the fungal secretome^62^. In both rabbit and murine models of *C. neoformans* infection, low-glucose conditions have been reported during early stages of infection, including lung colonization and intra-macrophage survival^63,64^. Consistent with these observations, we found that low glucose markedly altered EV composition and induced mitochondrial tubularisation across VNI isolates, with these effects most pronounced in the VNIa-5 subclade.

Our study demonstrates that glucose availability profoundly influences key virulent traits in *C. neoformans*, including EV composition and mitochondrial morphology. The fungus appears capable of dynamically modulating EV profiles in response to tissue glucose levels, potentially reshaping host immune responses and facilitating dissemination from lungs to the central nervous system^51^. In VNIa-5 strains, low-glucose conditions markedly increased mitochondrial tubularisation, a phenotype that may contribute to enhanced intracellular fitness and proliferation during CNS infection. In addition, mitochondrial tubularisation may confer increased resistance to oxidative stress and antifungal agents. Consistent with this, both *in vitro* low-glucose environments and brain-specific glucose milieu have been linked to increased tolerance to fluconazole and amphotericin B respectively^65,66^.

Together, these findings highlight the critical role of glucose adaptation as a central determinant of *C. neoformans* virulence. The clinical dominance of VNIa-5 in Asia may therefore reflect enhanced glucose responsiveness, manifested through dynamic modulation of EV composition and mitochondrial morphology. Nonetheless, as our conclusions are derived largely from genomic analyses and *in vitro* phenotypic assays, further *in vivo* studies will ultimately shed light on the ultimate contribution of these glucose-responsive traits to pathogenicity.

## Methods

### Strain collection and ethics statement

The 81 Chinese isolates included in this study consisted of 28 clinical isolates and 53 environmental isolates. Clinical isolates were collected from the cerebrospinal fluid (CSF) of HIV-negative patients at Shanghai Changzheng Hospital - a national reference hospital for cryptococcosis - between 2012 and 2023. All the participants involved gave written informed consent, and the study was approved by the Shanghai Changzheng Hospital Ethics Committee.

Environmental isolates were collected from pigeon droppings and tree hollows in urban areas across China. Decayed debris from tree hollows and pigeon droppings were sampled using cotton swabs and forceps, following previously described methods^7^. Samples were cultured on Niger seed agar, and the dark brown colonies were subjected to molecular identification via sequencing the internal transcribed spacer region. All isolates were stored in −80 °C freezer.

### Whole-genome sequencing and population genomics

To obtain whole-genome sequences, isolates were revived on Sabouraud dextrose agar (SDA) at 30 °C for 48 h. Single colony picks were then spread for confluent growth and incubated at 30°C for 24 h. Genomic DNA was extracted from approximately 0.5 g (wet weight) of yeast cells using the MasterPure Yeast DNA purification kit (Epicentre, USA) according to the manufacturer’s instructions. The genomic DNA pellet was resuspended in Tris-EDTA buffer and used to prepare paired-end sequencing libraries, then sequenced on HiSeq X10 platform or MGISEQ2000 platform (BGI-Wuhan, China) to deliver approximately 150 ⅹ coverage per strain. Paired-end reads were quality trimmed and then aligned by BWA-MEM (v 0.7.17) to the VNIa-5 reference genome assembly of strain BK42 generated using PacBio sequencing from Ashton’s study^6^. Funannotate (v1.8.16) was used for gene prediction and functional annotation of the VNIa-5 genome. Variants were then called using GATK (v 4.1.8.1) and filtered using “VariantFiltration” function with the parameters “QD < 2.0, FS > 60.0, MQ < 40.0”. Individual variants were filtered out if the minimum quality score was < 50, percent alternate allele < 0.8, or depth < 10.

Since VNIc has only been reported from Africa^4^, our population genomic analysis focused on VNIa and VNIb. For phylogenetic analysis of our 81 isolates, we included 8 previously sequenced *C. neoformans* isolates from Vietnam as reference^6^, representing each subclade of VNIa and VNIb (VNIa-5: BK42; VNIa-4: BK80; VNIa-93:04CN-03-050; VNIa-32: BK101; VNIa-outlier: BK158; VNIa-X:

BK230; VNIa-Y: NRHc5044; VNIb: BK8). A total of 155,594 variant sites were identified and concatenated for phylogenetic analysis. Phylogenetic trees were estimated using RAxML (v 8.2.12) under the “GTRCAT” model in rapid bootstrapping mode. For subdivision identification, all variant sites of the 89 sequences were concatenated and put into Admixture (v 1.3.0) with K running from 1 to 15. The cross-validation error value of each subdivision type was then compared to determine the optimized K value. Sliding-window population genetic statistics (π and F_ST_) were calculated with vcftools (v 0.1.16) per chromosome in 10k bp windows with “haploid” mode.

### Transcriptome analysis

Since transcriptome in YPD broth was reported to be more similar to *in vivo* cerebrospinal fluid (CSF) than in artificial CSF^67^, cells of three biological replicates were prepared by growing representative VNIa-5 strain (CH46) and VNIa-31 strain (CH14) separately in 30ml of YPD broth overnight at 37 °C. Total RNA was extracted by RiboPure-Yeast kit (Life Technologies, CA) and treated with DNase I (Life Technologies, CA) following the manufacturer’s instructions. The extracted total RNA was reverse transcribed and sequenced on a MGISEQ-2000 platform (BGI-Wuhan, China).

Raw RNA sequencing reads were filtered for adapter contamination and quality trimmed using SOAPnuke (http://github.com/BGI-flexlab/SOAPnuke), and reads containing adapters, or containing > 10% bases with quality < 10, or containing more than 1% unknown bases were discarded. Reads passing the filters were aligned to the annotated VNIa-5 genome using hisat (v 2.2.1) with parameters “–min-intronlen 20 –max-intronlen 1000”. Reads overlapping annotated transcripts of VNIa-5 were quantified with HTSeq-count (v 0.6.1) with matching mode set to “Union”. Read counts were normalized across samples and replicates using the Biocondutor package edgeR (v 4.0.14). Library sizes were normalized using trimmed mean of M values (TMM), a built-in algorithm in edgeR package. Differential expression was tested using the edgeR function “exactTest”. Significance values were corrected for multiple testing using the Benjamin-Hochberg false-discovery rate (FDR). Genes with an expression difference of at least 2-fold and an FDR *P* value below 0.001 were kept in the result. Enrichment analysis of the DEGs was conducted on DAVID website (https://davidbioinformatics.nih.gov/).

### EV isolation and nano particle tracking analysis (NTA)

EV isolation from solid agar media followed a recently published protocol^68^. Briefly, one colony of each isolate was inoculated into 5 mL of YPD medium and cultivated for 48 h at 30 °C with shaking. Solid medium for EV isolation under different glucose concentrations contained 6.7g yeast nitrogen base without amino acids (BD, USA), 1g Drop-Out mix (USBiological LifeSciences, USA), 0.4% ethanol, 3.3g NaCl, 2% agar, with high (2%) glucose or low (0.05%) glucose. The yeast cells were counted and diluted to a density of 10^7^ cells/ml in YPD. Aliquots of 300 μL of these cell suspensions were spread onto solid agar media and incubated for 24 h at 30 °C to reach confluence. Three petri dishes were used for each EV isolation. The cells were gently recovered from each of the three plates with an inoculation loop and transferred to a single centrifuge tube containing 30 mL of sterilized PBS. Suspended cells were centrifugation at 5,000 ⅹ g for 15 min at 4 °C. The cell pellet was resuspended in 1 mL PBS and OD_600nm_ was measured to determine the initial amount of the isolates. The supernatants were collected and centrifuged again at 15,000 ⅹ g for 15 min at 4 °C to remove debris. The resulting supernatants were filtered through 0.45 μm pore syringe filters and centrifuged at 100,000 ⅹ g for 1 h at 4 °C. Supernatants were discarded and pellets suspended in 300 μL of sterile PBS.

The amount and size distribution of EVs were measured using Zetaview (Particle Metrix, Germany) as described^69^. For this purpose, all samples were 200- to 1000- fold diluted in sterilized PBS and measured within the optimal dilution range of 1 to 9 ⅹ10^7^ particles/mL. Particle tracking analysis was performed in scatter mode with a 488 nm laser with the following settings: Sensitivity 70; shutter 70; minimum brightness 20; minimum area 5; minimum size 5; maximum size 1000, as suggested by the manufacturer. The EV amount was calibrated by initial yeast cell amount. The fraction of small EV or large EV was calculated by “EV_35-195nm_ sum amount/ total EV amount” or “EV_205-505nm_ sum amount/ total EV amount” respectively.

### Melanization measurement

Overnight cultures were harvested and washed twice with PBS at 3000 ⅹ g for 5 min for further phenotypic analysis. To prevent the influence of growth rate difference, we plated large number of cells on the plate initially. Thus, melanin production was assessed by plating 3 μL containing 10^6^ cells suspension on L-DOPA agar containing 1 g/L L-asparagine, 1 g/L glucose, 3 g/L KH2PO4, 250 mg/L MgSO_4_.7H_2_O, 1 mg/L Thiamine HCl, 5 μg/L Biotin, 100 mg/L L-DOPA, 20 g/L Bacto Agar. Plates were incubated in the dark at 30°C for 96 hours and imaged with same environment and light source. The figures were converted to gray scale image and the colony melanization level was measured using ImageJ software with “Invert” mode.

### *Cryptococcus* mitochondria morphology identification

Strains were grown and washed in PBS as described above and diluted to a final density of 10^6^ cells/mL in 10 mL of low-glucose (0.5 g/L) or high-glucose (4 g/L) DMEM (Beyotime biotechnology, China) in tissue culture flasks. DMEM with mild oxidative stress (0.7mM H2O2) for 24 hours is reported to be optimal condition for *in vitro* tubular mitochondria induction^46^ and was used for mitochondria observation under oxidative stress. After incubation for 24 hours at 37°C with 5% CO_2_, cells were again washed in PBS. To assay mitochondrial morphology, cells were then resuspended to 1 mL in PBS, stained for 1 hour at 37°C with 0.2 mM MitoTracker CMXRos (Beyotime biotechnology, China). After staining, cells were washed 3 times and resuspended in PBS. To quantify the fraction of cells with tubularized mitochondria, at least 100 yeast cells from 6 randomly chosen microscope fields were analyzed independently by two observers. The mitochondria morphology of cryptococcal isolates were observed and counted under 100 × oil immersion lens of Leica DM5000B fluorescent microscope. Images were collected on Olympus FV3000 laser scanning confocal microscope system with 60 × oil immersion lens.

### Macrophage infection and intracellular proliferation rate determination

The phagocytosis experiment followed the steps as previously described^47^. Briefly, 10^5^ J774 cells were plated into a 24-well plate 24 h before infection. One hour before infection, J774 cells were switched to low-glucose (0.5 g/L) DMEM and activated with 150 ng/mL phorbol 12-myristate 13-acetate. At the same time, 10^6^ yeast cells per 100 μL were opsonized with the monoclonal antibody 18B7 to cryptococcal capsule (Sigma, USA). After pre-incubation, the opsonized yeast cells were directly added to J774 cells at a ratio of ten yeast cells per macrophage and phagocytosis was allowed to proceed for 2 h. Non-internalized yeast cells were removed by 6-times washes with pre-warmed PBS.

For time point T=0, DMEM was removed and 200 μL of sterile distilled H2O was added into wells to lyse macrophage cells immediately after 2 h infection. After 30 min, the released intracellular yeasts were collected. Another 200 uL distilled H2O was added to each well to collect the remaining yeast cells. Intracellular yeasts were counted using plate CFU counting. For the subsequent time point (T=24h), intracellular cryptococcal cells were collected and independently counted. The intracellular proliferation rate (IPR) value was calculated by dividing the maximum intracellular yeast number by the initial intracellular yeast number at T=0.

### Dormant cell induction and intravenous mice infection

*In vitro* induction of viable but not cultivable (VBNC) / dormant yeast cells mainly followed the steps as previously described^70^. After the yeast had grown for 24 h, in which fungi reached the early stationary active phase (active stage 1), 100 μL of active stage 1 was inoculated into flask containing 10 mL of liquid YPD, which was incubated for 24 h, 150 RPM, at 30°C. After 24 h of growth, the fungus reached stationary growth again (active stage 2); we placed the flask in a hypoxia incubator (1% oxygen) and incubated it for 8 days at 30°C. Fungal cultivability and viability were monitored in each experiment to confirm the VBNC phenotype before proceeding to subsequent experiments.

For cultivability, the fungus was washed and diluted with PBS, and 10^4^ cells were plated on a YPD agar plate and incubated for 2 days at 30°C. After 2 days, the colonies were counted, representing the number of cultivable yeasts in relation to the initial number inoculated. Fungal cell viability was tested by Acridine Orange/Propidium Iodide (AO/PI) Double-Staining Assay (Beyotime biotechnology, China) according to the manufacturer’s recommendations (10 μL/mL for 10^6^ cells for 20 min). The incubated cells were then viewed under a Leica DM5000B fluorescence microscope at 20× lens with red or green channel. The images generated under the two channels were further merged using ImageJ software to examine viability of the strain.

The virulence of the dormant cells from three VNIa-5 and 3 VNIa-31 isolates was assessed using 7 ∼ 8-week-old female C57BL/6 mice with initial weight of 22-23g. Mice (6 per isolate) were inoculated intravenously with 100 μL of cell suspension (approximately 10^5^ dormant cells / mouse). The survival rate and the mice weights were observed until 40 days post-infection.

### Statistical analysis

Data was analyzed using R version 4.3.1. Continuous data sets were tested for normal distribution using the Shapiro-Wilks test and for homogeneity of variance using Levene Statistics, and if data sets fit the requirements, subjected to parametric analysis by *t*-test. If data sets were not normally distributed or failed to show homogeneity of variance, the non-parametric Mann-Whitney *U*-test was applied to test for statistically significant differences. Categorical data was analyzed by Fisher’s exact test. *P*-values < 0.05 after adjusting for multiplicity were considered statistically significant. Phenotypic experiments were performed with at least three repeating. Counting experiments to identify the fraction of yeasts with tubular mitochondrial morphology under microscope were performed in a blind manner. For *in vitro* phenotypic studies, yeast strains were randomly distributed in different culture well plate positions.

### Data availability

The sequencing data in this study have been deposited in the Genome Sequence Archive in National Genomics Data Center, China National Center for Bioinformation https://ngdc.cncb.ac.cn/gsa. The WGS data are under accession PRJCA044073 and RNAseq data are under accession PRJCA044471. The annotation data for VNIa-5 reference genome is available as a gff3 file from https://doi.org/10.6084/m9.figshare.29922578. All other data is available upon request.

### Code availability

Custom scripts for genomic and transcriptome analysis can be found at https://github.com/HongN/Cneo-analysis.

### Additional files

Table S1. Isolates information and MLST-based epidemiological analysis results.

Table S2. Genes located in VNIa-5 selection sweep regions. (“NA” Not available).

Table S3. Genes unique to VNIa-5.

Table S4. 381 DEGs between VNIa-5 and VNIa-31. (“NA” Not available).

Table S5. NTA results of cryptococcus extracellular vesicles.

Table S6. IPR values and mitochondrial tubularisation fraction

Table S7. Cultivability and viability of dormant *Cryptococcus neoformans*.

## Acknowledgments

This study was funded in part with the grants from National Natural Science Foundation of China (No. 82172291 and 82102419). M.C.F. received funding from the Wellcome Trust and the UK Medical Research Council MRC MR/R015600/; he is a member of the CIFAR Fungal Kingdoms program.

## Author contributions

H.N., B.X.X., Y.H.S. and Y.P. performed most experiments and bioinformatic analysis. H.E., M.Y.N., C.H.L., Z.Q., W.Y. and W.M.M. contributed to experiments and data analysis. L.W.Q. and C.M. provided critical clinical isolates. M.C.F., C.M., X.J.P. and H.N. supervised the study, performed data interpretation, and acquired funding. H.N. and M.C.F. wrote the paper with input from all co-authors. All authors read and approved the paper.

